# Attentional rhythms are generated in binocular cells

**DOI:** 10.1101/2022.04.05.487107

**Authors:** Bo Dong, Guangyao Zu, Jianrong Jia, Airui Chen, Ming Zhang

**Author notes:** Corresponding authors: Airui Chen, Ming Zhang. co-First author (Bo Dong, Guangyao Zu). **Contribution** Bo Dong: Conceptualization, Software, Data curation, Formal analysis, Funding acquisition, Investigation, Supervision, Project administration, Writing –original draft, Writing –review & editing. Guangyao Zu: Data curation, Formal analysis. Jianrong Jia: Formal analysis. Airui Chen: Conceptualization, Data curation, Formal analysis, Funding acquisition, Investigation, Supervision, Project administration, Writing –original draft, Writing –review & editing. Ming Zhang: Funding acquisition, Investigation, Supervision, Project administration.

## Abstract

Visual attention is intrinsically rhythmic and oscillates based on the discrete sampling of either single or multiple objects. Recently, studies employing transcranial magnetic stimulation (TMS) and high-temporal behavioral approaches have found that the early visual cortex (V1/V2) modulates attentional rhythms. However, both monocular cells and binocular cells are in the early visual cortex, and whether the neural site of attentional rhythms is monocular cells or binocular cells remains poorly understood. In Experiment 1, we reset the phase of attentional rhythms in one monocular channel (left eye or right eye) by the dichoptic cue and tracked the temporal response function (TRF) of the monocular channel in the left and right eyes separately using time response tracking technology. We found no significant differences in the two TRFs of each monocular eye, suggesting that attention rarely switched between the two eyes, indicating that binocular cells, not monocular cells, are the neural site of attentional rhythms. These results were verified even when resetting the phases of attentional rhythms by a binocular cue in Experiment 2. These results constitute direct neural evidence supporting rhythmic attention theory.

## 1 Introduction

Recently, some important studies found that attention systems work in a discrete rather than a continuous manner (Fiebelkorn, Saalmann, & Kastner, 2013; Landau & Fries, 2012; VanRullen, 2016). In the fine time dimension, visual attention manifests as a rhythmic spotlight and divides visual information into a series of temporal chunks with a length of several hundred milliseconds (Busch & VanRullen, 2010; Landau, Schreyer, Van Pelt, & Fries, 2015). This cyclical manifestation is called the attentional rhythm (Chen, Tang, Wang, & Zhang, 2017; Fiebelkorn & Kastner, 2019; Herbst & Landau, 2016; Jensen, Bonnefond, & VanRullen, 2012; VanRullen, 2016, 2018). Attentional rhythms modulate our perception performance and manifest in various behavioral oscillations. When people focus their attention on a single location or divide their attention to multiple locations (i.e., location-based attention), researchers have found theta and alpha oscillations in aspects of processing efficiency, such as the reaction time, accuracy, and sensitivity (d-prime) (Chen, Wang, Wang, Tang, & Zhang, 2017; Chen, Zu, Dong, & Zhang, 2020; Dugué, McLelland, Lajous, & VanRullen, 2015; Landau & Fries, 2012; Song, Meng, Chen, Zhou, & Luo, 2014; VanRullen, Reddy, & Koch, 2005; Zhang, Morrone, & Alais, 2019). Similarly, behavioral oscillations were found in object-based attention (Fiebelkorn et al., 2013) and feature-based attention (Mo et al., 2019; Re, Inbar, Richter, & Landau, 2019). Interestingly, some top-down and downtop factors, such as the saliency of the target, predictability of cues, task difficulty, task rewards and cortical distance, could modulate the patterns of attentional rhythms (Chen, Wang, et al., 2017; Chen et al., 2020; Dugué, Xue, & Carrasco, 2017; Jia, Liu, Fang, & Luo, 2017; Su, Wang, Kang, & Zhou, 2021). Although attentional rhythms are robust and evident in different paradigms of visual attention, our knowledge regarding the neural site(s) of this phenomenon and how our brain controls the temporal characteristics of attention is limited (VanRullen, 2016).

The early visual cortex (V1/V2) may be involved in the production of attentional rhythms. The early visual cortex is the first neural site of visual information processing in the cerebral cortex, and its role in spatial attention has been confirmed by numerous studies (Hembrook-Short, Mock, Martin Usrey, & Briggs, 2019; Klein et al., 2018; Motter, 1993). Dugué et al. (2015) used the transcranial magnetic stimulation (TMS) technique to conduct research related to this topic and found that compared with the condition of no TMS interference, search efficiency was significantly reduced when TMS was applied over the occipital cortex (V1/V2). In addition, these researchers found that the degree of reduction alignment to the temporal moment of applying TMS pulses showed periodic fluctuations, i.e., behavioral oscillations. More importantly, when the applied frequency of TMS is close to the frequency of behavioral oscillations (i.e., 5.7 Hz), the decrease in search efficiency in the visual search task (Dugué, Marque, & VanRullen, 2015) and the detection accuracy in the cue-target task (Dugué, Roberts, & Carrasco, 2016) were the largest, suggesting that interference with V1/V2 can significantly affect behavioral oscillations and indicating the role of V1/V2 in attentional rhythms (Dugué & Van Rullen, 2017). However, TMS stimuli generate interfering currents in the cerebral cortex through a rapidly changing magnetic field. Even if a high-precision type of coil is used, the radius of the interfering area is approximately 2∼5 cm (Wassermann, 2008), and it is impossible to accurately determine where and the extent to which TMS stimuli interfere with V1/V2. In addition, V1/V2 is a complex cortical area (Wandell, 1995) that contains both monocular and binocular vision pathways (Hubel & Wiesel, 1977). Hence, the visual pathway impacted by TMS pulses to generate the effects described above remains unclear.

According to the anatomical characteristics of the visual pathway, monocular visual information in mammals and humans is first transmitted to the left and right visual cortex separately. Only when information is transmitted to the fourth layer of cells (IVc) in V1 can the information from both eyes converge. In this precortex pathway (e.g., retina and lateral geniculate nucleus), the visual signals from one eye are independent of the visual signals from the other eye, and each monocular cell can only receive information from the corresponding eye (same-eye condition) and cannot receive information from the opposite eye (different-eye condition); thus, monocular cells can provide information about the eye of origin (Livingstone & Hubel, 1987). These monocular cells can also be found in V1 and constitute the ocular-dominant columns. As these visual signals continue to forward (e.g., layers II and III in V1, V2, V3, etc.), the separate signals from the opposite eyes can converge to a single cell (i.e., binocular cells) (Hubel & Wiesel, 1977). The signal from either the left or right eye can activate the binocular cell, while the information about the eye of origin disappears. According to the anatomical characteristics of the visual pathway, researchers can apply the following logic: if a visual phenomenon differs between same-eye and different-eye conditions, the neural site of the phenomenon is located before the IVc layer of V1, i.e., the monocular visual pathway; otherwise, if there is no difference between same-eye and different-eye conditions, the neural site of the phenomenon is generated in the binocular visual pathway, i.e., the IVc layer of V1 and other higher cortical areas (V2, V4, IT, or LIP). According to this logic, Zhaoping (2008) found that the saliency map is produced by V1 neurons. The study also found that compared to the target and distracters appearing in the same eye, more time was required for the participants to find the target when the target and distracters appeared in the visual field of different eyes. This discrepancy indicates that the attention saliency map is created during the monocular cell processing stage in V1 (Zhaoping, 2008). In addition, the logic of comparing individuals’ behavioral or neural performances between same-eye and different-eye conditions has been widely used in the investigation of the neural sites of other visual perception phenomena, such as orientation adaptation (Gilinsky & Doherty, 1969), spatial frequency adaptation (Blakemore & Campbell, 1969), color adaptation (McCollough, 1965), motion aftereffect (Anstis, Verstraten, & Mather, 1998), subjective contour perception (Paradiso, Shimojo, & Nakayama, 1989), and perceptual learning (Schoups & Orban, 1996).

The unique anatomical structure of V1 may be used to explore the neural sites of attentional rhythms. Pertinently, in a recent study, we adopted a high-time-resolution cue-target task in combination with a stereoscope to control the presentation of cues and targets presented in the visual fields of the left and right eyes. The pattern of behavioral oscillations when the cue and target were presented in the visual field of the same single eye was similar to that under the different single-eye condition. Thus, the presentation in the visual field of the same or different eyes did not affect the behavioral oscillations; thus, this study indicates that the neural site of attentional rhythms is located in the binocular cell visual pathway, where information about the eye of origin is lost (Chen, Wang, Wang, Tang, & Zhang, 2018). However, notably, in this study, the cue and target were randomly presented in the following four areas: the left eye and left visual field, the left eye and right visual field, the right eye and left visual field, and the right eye and right visual field (see Chen et al. (2018) in Figure 2). This manipulation resulted in two conditions, namely, a cue-target condition in the same eye and a cue-target condition in different eyes. Thus, in the experiment, the cue and target switched randomly between not only the left and right eyes but also the left and right half of the visual field. Notably, the attentional system not only oscillates between the left and right eyes (interocular oscillation) but also switches between the left and right visual fields (visual field oscillation). Interocular oscillations and visual field oscillations occur simultaneously, and it is difficult to separate the effects of the two types of oscillations from behavioral fluctuations. To eliminate the interference of visual field oscillations, we only presented monocular stimuli in the central visual field of the left and right eyes to investigate the neural site of attentional rhythms.

The temporal response function (TRF) is an effective method used to measure neural activity underlying attentional rhythms (Jia, Fang, & Luo, 2019; Jia et al., 2017; Vanrullen & MacDonald, 2012). This method has been successfully used to study the temporal dynamics of working memory (Huang, Jia, Han, & Luo, 2018) and visual crowding (Han & Luo, 2019). This method assumes that the visual system is linear and that the neuron response at each moment is the convolution of the neural activity induced by all previous stimuli. Using this approach, we can record the electrical signals of the brain (i.e., the output of the linear system) and simultaneously present a sequence of visual stimuli whose basic feature characteristics (such as brightness, contrast, and movement direction) are rapidly and randomly changed (i.e., the output of the linear system), and the brain electrical signal (i.e., the output of the linear system) is recorded. By performing deconvolution calculations based on the input and output, the TRF of the visual system for this feature can be obtained (Crosse, Di Liberto, Bednar, & Lalor, 2016; Lalor, Pearlmutter, Reilly, McDarby, & Foxe, 2006). The TRF represents the response patterns of the brain (system) to a certain stimulus (input). The TRF characterizes classical components, such as C1, P1, and N1, in event-related potential (ERP) research and is not affected by psychological processes, such as semantics and emotions. In addition, researchers can manipulate various sequences of feature characteristics to track neural responses to multiple locations, features, or objects simultaneously, which is similar to the steady-state visual evoked potential (SSVEP) technique. Therefore, time response tracking technology is an effective method for investigating attentional rhythms. Furthermore, we can obtain a single TRF or multiple TRFs within 0.5 to 1 hour, which is less time than that needed for the high-temporal behavioral method to measure attentional rhythms. This saving in time can not only better ensure the stability of the participants’ attentional state during the experiment but can also decrease the time needed and cost of the study. In the classic cue-target paradigm, the TRFs of the left stimulus and the right stimulus are calculated using time response tracking technology, and the antiphase pattern of these two TRFs is revealed (Jia et al., 2019; Jia et al., 2017). More importantly, the significant frequencies of the neural oscillations are consistent with the behavioral oscillations, indicating that TRF is an effective measurement index for attentional rhythms (Jia et al., 2019; Jia et al., 2017).

To observe the neural activity of attentional rhythms, this study adopted temporal response tracking technology to calculate the TRFs of stimuli presented in the visual fields of participants’ left and right eyes. We aimed to examine whether an antiphase or in-phase relationship exists between the two TRFs. Then, we identified whether the attentional system oscillates between the two eyes and investigated the role of V1 in attentional rhythms. Based on our work (2018), in Experiment 1, we presented stimuli in the central visual field of each eye to eliminate the confounding effects of oscillations between visual fields and simultaneously used temporal response tracking technology to obtain the two TRFs of the brain when processing monocular stimuli projected into the visual field of each eye. If the neural sites that generate attentional rhythms are located in the monocular visual pathway, a significant difference between the TRFs under the cued-eye condition and uncued-eye condition and an obvious antiphase pattern of neural activity should be observed. In contrast, if the neural sites are located in the binocular pathway, the TRFs under the two conditions should be in the same phase, and there should be no significant difference. To further confirm that the phase result of the TRFs between the two eyes in Experiment 1 was caused by oscillations between the visual fields of the two eyes, we conducted a control experiment (Experiment 2). We made the attentional system oscillate synchronously in the two eyes by cuing both eyes simultaneously to eliminate visual field oscillations and interocular oscillations. Thus, we sought to obtain two TRFs when the attentional rhythms in both eyes were exactly in the same phase to verify the speculation from Experiment 1.

## 2 Experiment 1: Attentional rhythms under the monocular cuing condition

### 2.1 Participants

Ten participants (8 females and 2 males, all right-handed) were recruited for Experiment 1 and provided written consent. The participants were aged 19 to 23 years (*M* = 21.1 years, *SD* = 1.45 years). All participants had normal or corrected-to-normal visual acuity, no color blindness, and no color weakness. The participants were paid after the experiment. This study was conducted in accordance with the Declaration of Helsinki and was approved by the ethical committee of Suzhou University of Science and Technology.

#### 2.1.2 Design

A single factor with a two-level within-subject experimental design was used to investigate the interactive patterns of attentional rhythms in the left and right eyes of the participants. According to blinking spotlight theory, the attentional system always rhythmically oscillates, which enables our attention to discretely process a single object and periodically switch between multiple objects (Vanrullen, 2013, 2016, 2018). The phases of attentional rhythms are random and disorganized at different time points in each individual. To compare the phases of the TRF of the left and right eyes, we presented cue stimuli in the visual field of one eye to reset the chaotic phases of attentional rhythms; this manipulation is similar to that used in previous studies (Fiebelkorn et al., 2013; Jia et al., 2019; Jia et al., 2017; Landau & Fries, 2012). Then, we observed and measured the phase relationship between the TRFs in the cued eye and the uncued eye. If an obvious antiphase relationship was present, the attentional system should be able to oscillate between the two eyes rather than only object(s). Pertinently, the neural sites of attentional rhythms are located in monocular cells. Otherwise, attentional rhythms originate from neural sites in binocular cells.

To obtain the two TRFs, one for the cued eye and one for the uncued eye, we used two randomly independent sequences to control the contrast of each disc presented in the visual field of each eye (Jia et al., 2017). Similar to the SSVEP technique, temporal response tracking technology can trace and separate neural activity in response to stimuli in both cued and uncued eyes simultaneously in each trial. Importantly, the TRF represents a left- or right-eye-specific tracking response and does not depend on the presence or absence of the target. To maintain the participant’s attention state in the experiment, the target was presented in only 25% of the trials.

#### 2.1.3 Apparatus

The procedures were written by MATLAB and Psychophysics Toolbox-3 (Brainard, 1997; Pelli, 1997). The experiment was run on a Dell XPS8700 computer equipped with a GTX1050Ti graphics card. The display was a 22-inch ViewSonic P225f CRT with a resolution of 1024×768 and a refresh rate of 60 Hz. A keyboard was used to document the responses. After linear correction of the screen, a Bits# device was used to generate the stimuli to ensure the temporal precision of the presentation of the visual stimuli. Furthermore, we adopted Bits# to send triggers to the NeuroScan system to ensure the synchronization precision of the EEG signal and stimuli presentation. Stereoscopes were used to separately reflect images from both sides of the computer screen onto the participants’ field of view and the corresponding eyes.

#### 2.1.4 Stimuli

All stimuli were presented on a gray (7.53 cd/m^2^) background. The outer diameter of the two-disc stimuli was 8°, and the inner diameter was 1°. The two discs were black and white checkboard patterns. Using stereoscopes, the left and right images in the central visual field of both eyes were fused, and the participants could see only the discs. The participants were required to focus on the fixation cross (horizontal and vertical 1° visual angle), which was in the center of the disc stimulus. To improve the fusion of the left and right images, we presented one high-contrast black and white ring (width: 0.67° visual angle) at a 2° visual angle from the outside of the discs and 12 other high-contrast black and white rings (width: 0.17° visual angle) in the center of the discs. We generated two temporal sequences with equal power at all frequencies after inverse FFT. The contrast of the two discs was modulated according to two independent sequences in time aligned to the refreshed frame, i.e., spread spectrum stimuli (Crosse et al., 2016). The luminance of the screen was set from 0.01 cd/m^2^ (black) to 67.96cd/m^2^ (white). The cue was a white ring (width: 1° visual angle) that was presented in the cued eye for 500 ms. The target event was a contrast decrement of 500-ms duration. The contrast of the target was computed by the Staircase procedure so that the task difficulty of the experiment could be set at a 79.4% threshold. The target appeared randomly on a certain area of some disc but avoided overlapping with the central fixation point.

#### 2.1.5 Procedures

The experiment was conducted in a dark room. The participants sat in front of a computer 68 cm away from the screen, with their heads stabilized on a chin rest. Each participant was allowed to adjust the stereoscope until the images projected into the visual fields of the left and right eyes were fused well. In each trial, the static disc stimuli were presented for 3000 ms, and a white circle with a width of 1° appeared around the disc stimulus as the cue in a single eye. After the cue disappeared, the two discs began to flicker and continued to flicker for 5000 ms. In 25% of the trials, while the discs flickered (250 ms to 4250 ms), the target appeared randomly on a certain disc in the visual field of the cued eye or uncued eye. After the discs disappeared, a white square (with a side length of 1°) appeared at the fixation point, and the participants were asked to press a key. The participants needed to determine whether a target was present in each trial. If the target was present, the participants pressed the left arrow on the keyboard. Otherwise, the participants pressed the right arrow on the keyboard. After their responses were collected, after each trial, feedback was presented on the screen, i.e., correct, incorrect, or no response. The participants completed the tasks as accurately as possible. There were 128 trials distributed equally across 6 sessions. The participants could rest after each session.

### 2.2 Data analysis

#### 2.2.1 EEG recording and preprocessing

The EEG data were collected using Synamps 2 amplifiers equipped with a 10-20 standard extended 64-channel ActiCap (Neuroscan). The sampling rate was 1000 Hz. The data were grounded with the electrode placed in front of the forehead and referenced to location Fz. The vertical electrooculography (EOG) signals were recorded by two electrodes placed above and below the left eye, and the horizontal EOG signals were recorded by another two electrodes placed around the left and right eyes. During the experiment, the impedance of the electrodes was kept below 5 kΩ.

According to previous studies (Jia et al., 2019; Jia et al., 2017), we used the FieldTrip toolbox to process the EEG data. The trials were rereferenced to the average of all electrodes, bandpass filtered from 2 to 50 Hz, and then epoched from 0 to 5000 ms alignment to the onset of the disc flickering using a 500 ms prestimulus baseline correction. To eliminate the effects of onset and offset responses activated by the flicking stimuli on computing the TRF, the stimulus time points in the 3.5 s before the last 1 s and after 0.5 s of the sequence were entered in the TRF estimation. Before the TRF was computed, the EEG data and sequences of contrast were spliced together sequentially and converted into z scores.

#### 2.2.2 Calculation of TRFs

We calculated the TRFs using two contrast sequences of stimuli and the corresponding EEG data. The convolution function between the sequence and the corresponding EEG data is R(t) = TRF*S(t). R(t) represents the neural response, S(t) represents the stimulus input, and * represents the convolution process. According to this formula, the disc contrast sequence and EEG data are deconvolved to calculate the TRF of the visual system in response to the disc stimulus contrast. The parameter lambda is used to control overfitting in ridge regression, and the lambda parameter of all participants was set to 1 (Jia et al., 2019; Jia et al., 2017). Then, we calculated the cued-eye and uncued-eye TRFs for each electrode and each participant. Next, according to previous studies and the topographic map of the EEG data, the data from electrodes POz and Oz were selected for further analysis.

#### 2.2.3 Frequency and phase analysis

Fast Fourier transform (FFT) was used to analyze the frequency of the TRFs of the cued eye and uncued eye for each participant, and a permutation test was adopted to measure the significance of the frequencies (Chen, Wang, et al., 2017; Chen et al., 2020; Landau & Fries, 2012). After the TRF was calculated, the original value of the TRF was randomly shuffled; thus, the corresponding map between the original TRFs and time points was shuffled. In addition, an FFT analysis was performed to calculate the energy that may be generated by random factors in each frequency of the TRF. We shuffled 1000 times to obtain the amplitude corresponding to 1000 pseudosignals at each frequency. These amplitudes constitute the permutation distribution of each frequency. By calculating the percentage of the original energy at each frequency, the significant oscillation patterns were obtained. Since multiple comparisons were involved in the permutation test, the false discovery rate (FDR) method was used to correct for multiple comparisons.

### 2.3 Results

In Experiment 1, we presented a cue in the visual field of a single eye in combination with a TRF approach to examine the corresponding TRF responses in the cued eye and uncued eye and investigate the role of monocular and binocular cells in attentional rhythms. The Oz and POz electrodes were close to the early visual cortex and are usually selected for analysis in the TRF approach (Lalor et al., 2006). Therefore, we selected these two electrodes for the frequency and phase analysis. Clear, stable, and effective TRF components are prerequisites for investigating attentional rhythms. To determine the credibility of the TRF response, we shuffled the temporal sequence of the contrast, i.e., the convolution relationship in the linear system was shuffled, and the pseudo-TRFs induced by random factors were obtained (see the dotted line in Figure 2A and Figure 3A). Compared with the pseudo-TRFs, the TRF response contains obvious C1, P1, and N1 components, which is consistent with the findings of previous studies (Jia et al., 2017; Lalor et al., 2006). This finding suggests that the TRF approach in this study was credible and that the TRF response can be used as a neural indicator of attentional rhythms in further analysis.

#### 2.3.1 Oz electrode

We used an FFT analysis and permutation tests to examine the neural activity recorded at electrode Oz. The TRF responses showed strong theta band (2.7-6.3 Hz) activation under the cued-eye condition and theta-band (3.1-5.1 Hz) and alpha-band (10-11 Hz) activations under the uncued-eye condition (*p* < 0.05, FDR corrected). The findings regarding these two bands are consistent with the results of behavioral oscillation research (Dugué, Marque, et al., 2015; Fiebelkorn et al., 2013; Landau & Fries, 2012). This finding suggests that attentional rhythms were obvious under the cued-eye and uncued-eye conditions (see Figure 2B). To compare the phase differences in the TRFs under the cued-eye and uncued-eye conditions, we selected two wide ranges of significant frequencies (2.7-6.3 Hz and 10∼11 Hz) for the two TRFs. Figure 2C shows the phase relationship by the phase analysis. The phase difference in the 2.7-6.3 Hz frequency band between the TRFs under the cued-eye and the uncued eye conditions was 23.72° (Rayleigh test, *Z* = 5.64, *p* = 0.002), which showed no significant difference from 0° (i.e., in-phase mode) and significantly differed from 180° (i.e., antiphase mode; *μ* = 23.72°, 95% CI = [-7.61°, 55.06°]). The phase difference in the 10–11 Hz frequency band between the TRFs under the cued-eye and the uncued eye conditions was 5.74° (Rayleigh test, *Z* = 2.42, *p* = 0.087), which showed no significant difference from 0° (i.e., in-phase mode) and significantly differed from 180° (i.e., antiphase mode; *μ* = 5.74°, 95% CI = [-55.79°, 67.28°]). This finding suggests that the two TRFs show significant oscillation patterns; however, the two TRFs under the cued-eye and the uncued-eye conditions depict an in-phase mode instead of an antiphase mode. Notably, we also conducted other analyses, such as selecting a narrower range (such as 3.1∼5.1 Hz) and performing a point-by-point phase analysis, and we found the same phase relationship using these two analyses as the wider range analysis.

#### 2.3.2 POz electrode

We analyzed the neural activity recorded at electrode POz. The TRF responses showed strong theta band (2.7–5.5 Hz) activation and alpha-band (10-11 Hz) activation under the cued-eye condition and theta-band (3.1–5.1 Hz) activation under the uncued-eye condition (*p* < 0.05, FDR corrected). This finding suggests that attentional rhythms were also obvious under the cued-eye and uncued-eye conditions (see Figure 3B). Figure 3C shows the phase relationship. The phase difference in the 2.7–5.5 Hz frequency band between the TRFs under the cued-eye and the uncued-eye conditions was 1.84° (Rayleigh test, *Z* = 3.23, *p* = 0.036), which showed no significant difference from 0° (i.e., in-phase mode) and significantly differed from 180° (i.e., antiphase mode; *μ* = 1.84°, 95% CI = [-46.18°, 49.85°]). The phase difference in the 10–11 Hz frequency band between the TRFs for the cued-eye and the uncued-eye was -0.12° (Rayleigh test, *Z* = 2.33, *p* = 0.095), which showed no significant difference from 0° (i.e., in-phase mode) and significantly differed from 180° (i.e., antiphase mode; *μ* = -0.12°, 95% CI = [-63.96°, 63.71°]). This outcome suggests that the two TRFs show significant oscillation patterns; however, the two TRFs under the cued-eye and the uncued-eye conditions depict an in-phase mode instead of an antiphase mode.

By combining the frequency and phase results of the TRFs at the Oz and POz electrodes, resetting the phase of attentional rhythms in a single eye was not observed to induce the antiphase mode of the TRFs in two eyes; however, a stable in-phase mode was observed. This finding suggests that the attentional system does not oscillate back and forth between the left eye and right eye. In contrast, a synchronized oscillation mode between the two eyes is evident, indicating that attentional rhythms originate from binocular cells in the visual pathway of V1 and the higher visual cortex rather than monocular cells, which is consistent with our previous behavioral oscillation research (Chen et al., 2018). Notably, although the in-phase oscillation mode induced by monocular cues can rule out the possibility that monocular cells in the visual pathway produce attentional rhythms, whether the TRFs produced by a monocular cue (Experiment 1) are the same as the TRFs when binocular cells are activated is still unknown.

## 3 Experiment 2: Attentional rhythms under binocular cuing conditions

To further investigate the credibility of the TRF in-phase oscillation in Experiment 1, in Experiment 2, the cue was presented in the left eye and right eye simultaneously to reset the phases of the attentional rhythms in both eyes; this artificially made the attention system oscillate synchronously between the two eyes. If the interaction mode of the TRFs in both eyes observed in Experiment 2 is the same as that in Experiment 1, the results in Experiment 1 are suggested to be caused by “attentional synchronization oscillation”, thus indicating that attentional rhythms originate from binocular cells, which can be activated by both monocular cues (Experiment 1) and a binocular cue (Experiment 2).

### 3.1 Methods

Except for the cue stimulus in Experiment 2 being presented in both eyes (see Figure 1) simultaneously, the other stimuli and analysis methods were the same as those in Experiment 1.

**Figure 1.**
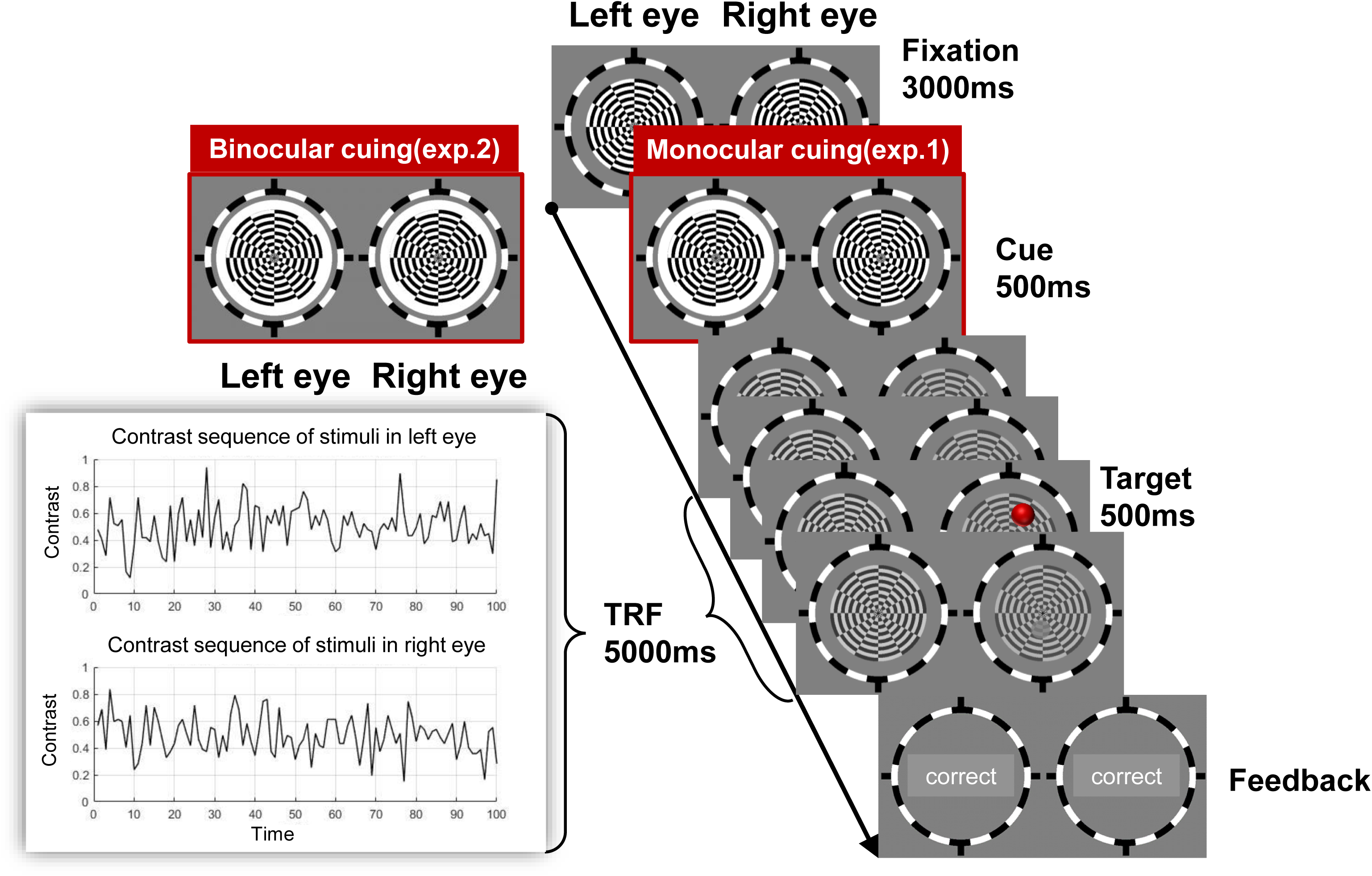
The procedures of Experiment 1 and Experiment 2. The red target in the figure is only for illustration. In the actuacxperiment, the target was an unobvious transparency change in a circular area.

**Figure 2.**
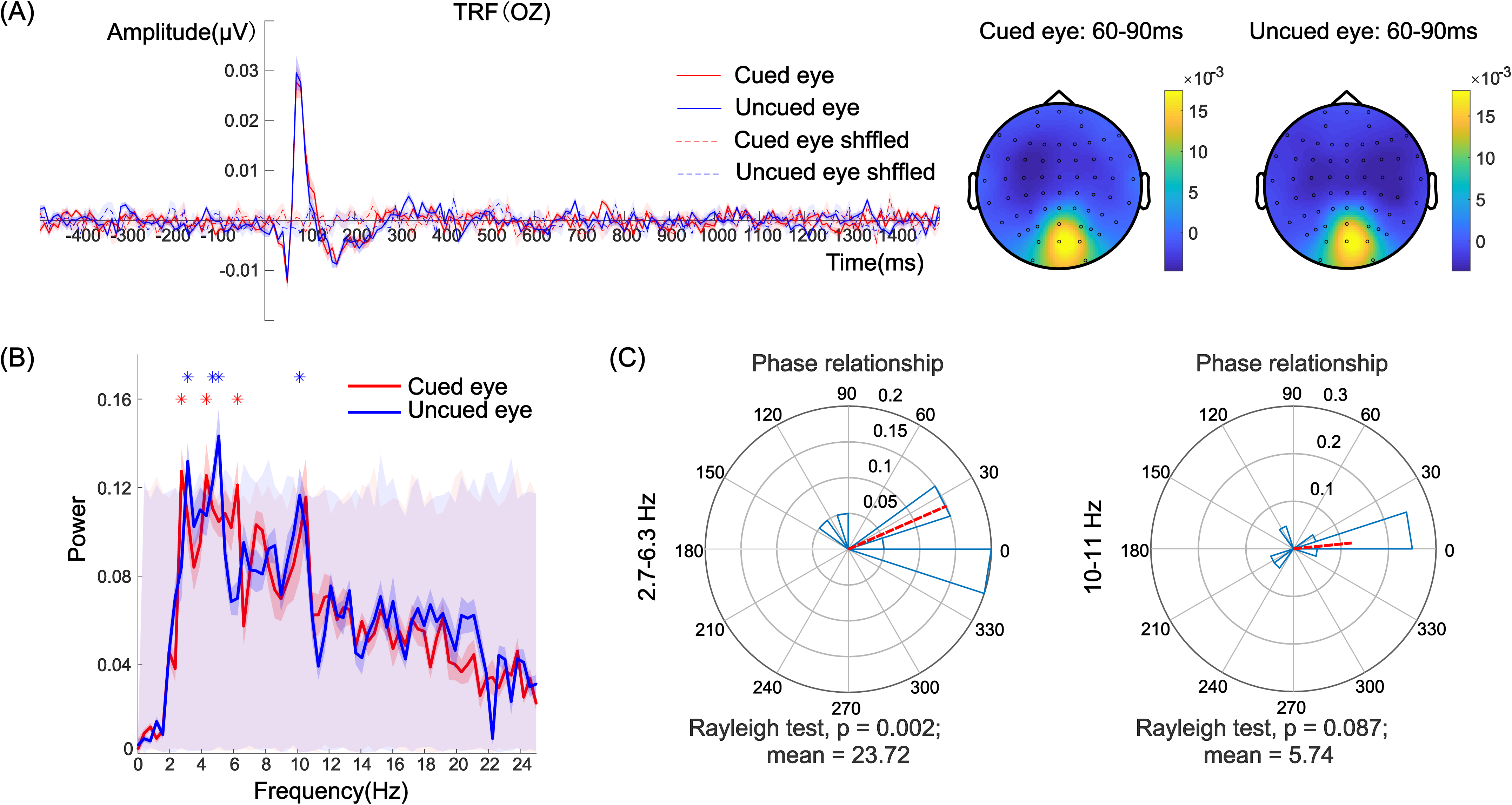
(A) TRF responses and topographic map under the cued-eye and uncued-eye conditions at electrode Oz. (B) Frequency map of the TRF waveform under the cued-eye and uncued-eye conditions. The asterisk represents significant frequency points after the permutation test under each condition. The gray shaded area with red low contrast represents the frequency range of 1000 pseudo-TRFs. (C) Phase relationship between the cued-eye and uncued-eye conditions with 2.7-6.3 Hz (theta band) and 10–11 Hz (alpha band) neural activity.

**Figure 3.**
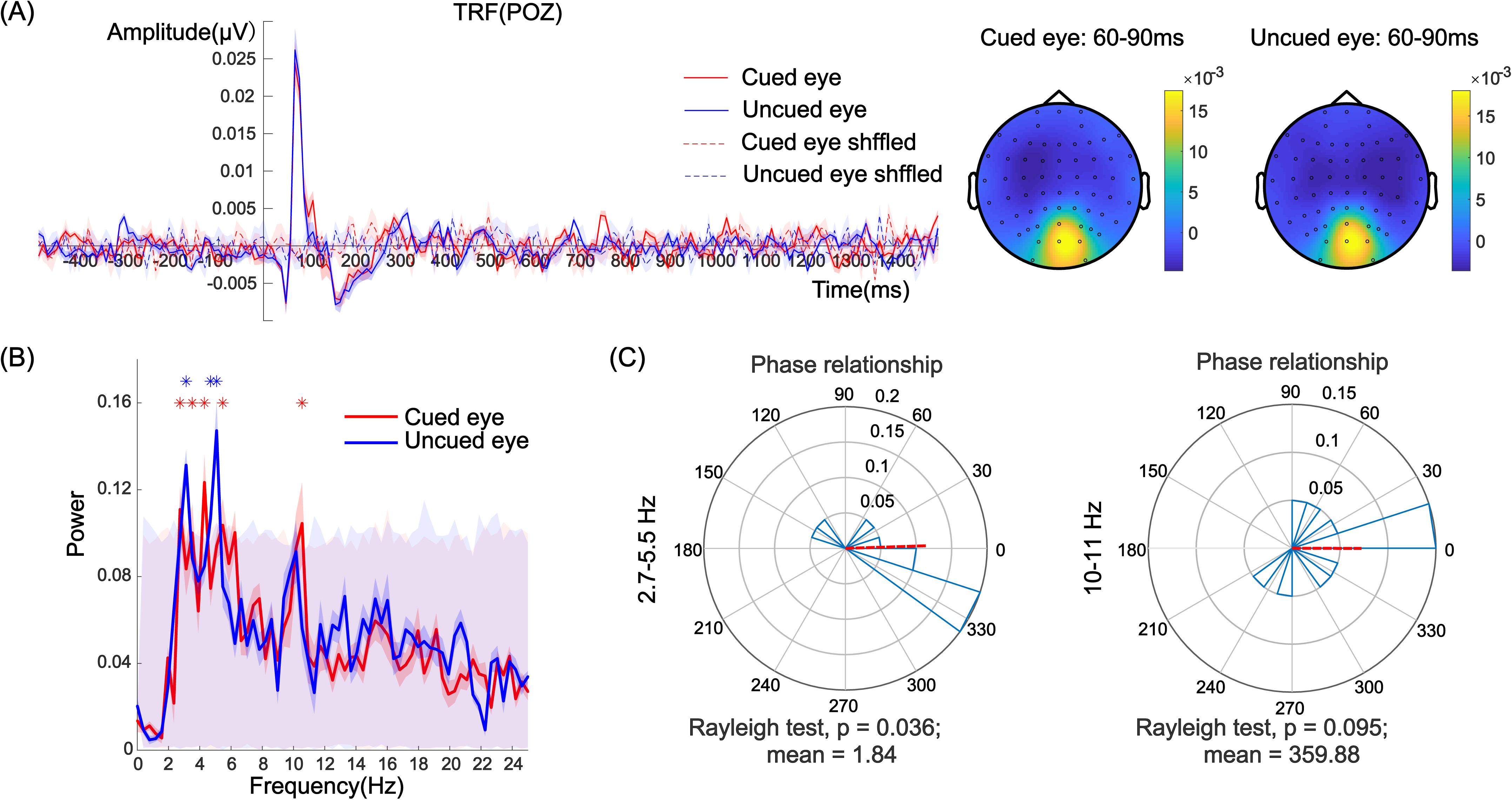
(A) TRF responses and topographic map under the cued-eye and uncued-eye conditions at electrode POz. (B) Frequency map of the TRF waveform under the cued-eye and uncued-eye conditions. The asterisk represents significant frequency points after the permutation test under each condition. The gray shaded area with red low contrast represents the frequency range of 1000 pseudo-TRFs. (C) Phase relationship between the cued-eye and uncued-eye conditions with 2.7-5.5 Hz (theta band) and 10-11 Hz (alpha band) neural activity.

### 3.2 Results

In Experiment 2, we aimed to explore the interaction mode of the TRF responses between the left eye and right eye when the cue was presented in both eyes simultaneously. Because the cue stimulus was presented in the left and right eyes simultaneously, we extracted the neural responses corresponding to the contrast sequences of the stimuli in the left and right eyes instead of those corresponding to the cued eye and uncued eye. To determine the credibility of the TRF waveform, we also shuffled the temporal sequences of the stimulus contrast to obtain the pseudo-TRF induced by random factors (see Figure 4A and the dashed part). Compared with the pseudo-TRF, the TRF signal in Experiment 2 contained obvious C1, P1, and N1 components, which is consistent with previous studies (Jia et al., 2017; Lalor et al., 2006), thus indicating that the TRF in this experiment is highly reliable. Therefore, the TRF can be used as a physiological indicator of attentional rhythms in further analysis.

**Figure 4.**
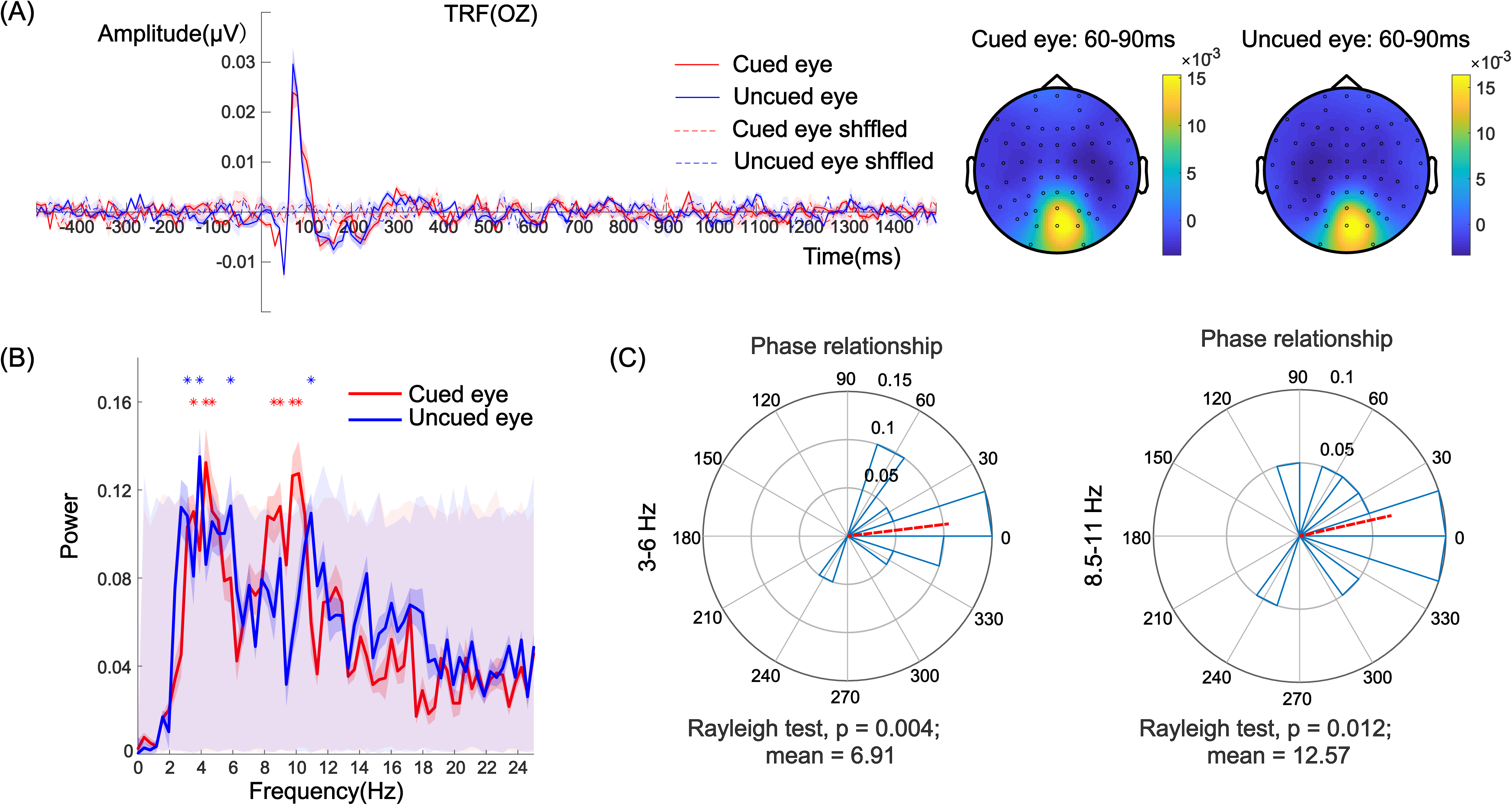
(A) TRF responses and topographic map of the two eyes at electrode Oz. (B) Frequency map of the TRF waveform of the two eyes. The asterisk represents significant frequency points after the permutation test under each condition. The gray shaded area with red low contrast represents the frequency range of 1000 pseudo-TRFs. (C) Phase relationship under 3-6 Hz (theta band) and 8.5-11 Hz (alpha band) neural activity.

#### 3.2.1 Oz electrode

We used an FFT analysis and permutation tests to examine the neural activity recorded at electrode Oz. The TRF responses showed strong theta band (3.5–4.7 Hz) and alpha-band (8.5–10.2 Hz) activations in the left eye and theta-band (3-6 Hz) and alpha-band (10-11 Hz) activations in the right eye (*p* < 0.05, FDR corrected). The findings of these two bands are consistent with the results in Experiment 1. This finding suggests that attentional rhythms were obvious under each eye condition (see Figure 4B). To compare the phase differences in the TRFs of the two eyes, we conducted a phase analysis in two ranges of frequencies (see Figure 4C). The phase difference in the 3–6 Hz frequency band in the TRF responses between the two eyes was 6.91° (Rayleigh test, *Z* = 4.98, *p* = 0.004), which showed no significant difference from 0° (i.e., in-phase mode) and significantly differed from 180° (i.e., antiphase mode; *μ* = 6.91°, 95% CI = [-27.59°, 41.41°]). The phase difference in the 8.5–11 Hz frequency band in the TRFs was 12.57° (Rayleigh test, *Z* = 4.14, *p* = 0.012), which showed no significant difference from 0° (i.e., in-phase mode) and significantly differed from 180° (i.e., antiphase mode; *μ* = 12.57°, 95% CI = [-27.08°, 52.21°]). This finding suggests that the two TRFs showed significant oscillation patterns. More importantly, the TRF responses portrayed an in-phase mode instead of an antiphase mode.

#### 3.2.2 POz electrode

We analyzed the neural activity recorded at electrode POz. The TRF responses showed strong theta band (3–5.5 Hz) and alpha-band (8.9–10.2 Hz) activations in the left eye and theta-band (3.9–6 Hz) activation in the right eye (*p* < 0.05, FDR corrected). The findings of these two bands are consistent with the results of Experiment 1. This finding suggests that attentional rhythms were obvious under each eye condition (see Figure 5B). To compare the phase differences in the TRFs between the two eyes, we conducted a phase analysis in two ranges of frequencies (see Figure 5C). The phase difference in the 3–6 Hz frequency band in the TRF responses between the two eyes was 19.34° (Rayleigh test, *Z* = 4.28, *p* = 0.01), which showed no significant differences from 0° (i.e., in-phase mode) and significantly differed from 180° (i.e., antiphase mode; *μ* = 19.34°, 95% CI = [-19.29°, 57.98°]). In addition, the neural activity of the 8.9–10.2 Hz frequency band showed no significant phase relationship (Rayleigh test, *Z* = 1.80, *p* = 0.17). This finding suggests that the two TRFs show significant oscillation patterns with an in-phase relationship instead of an antiphase relationship.

**Figure 5.**
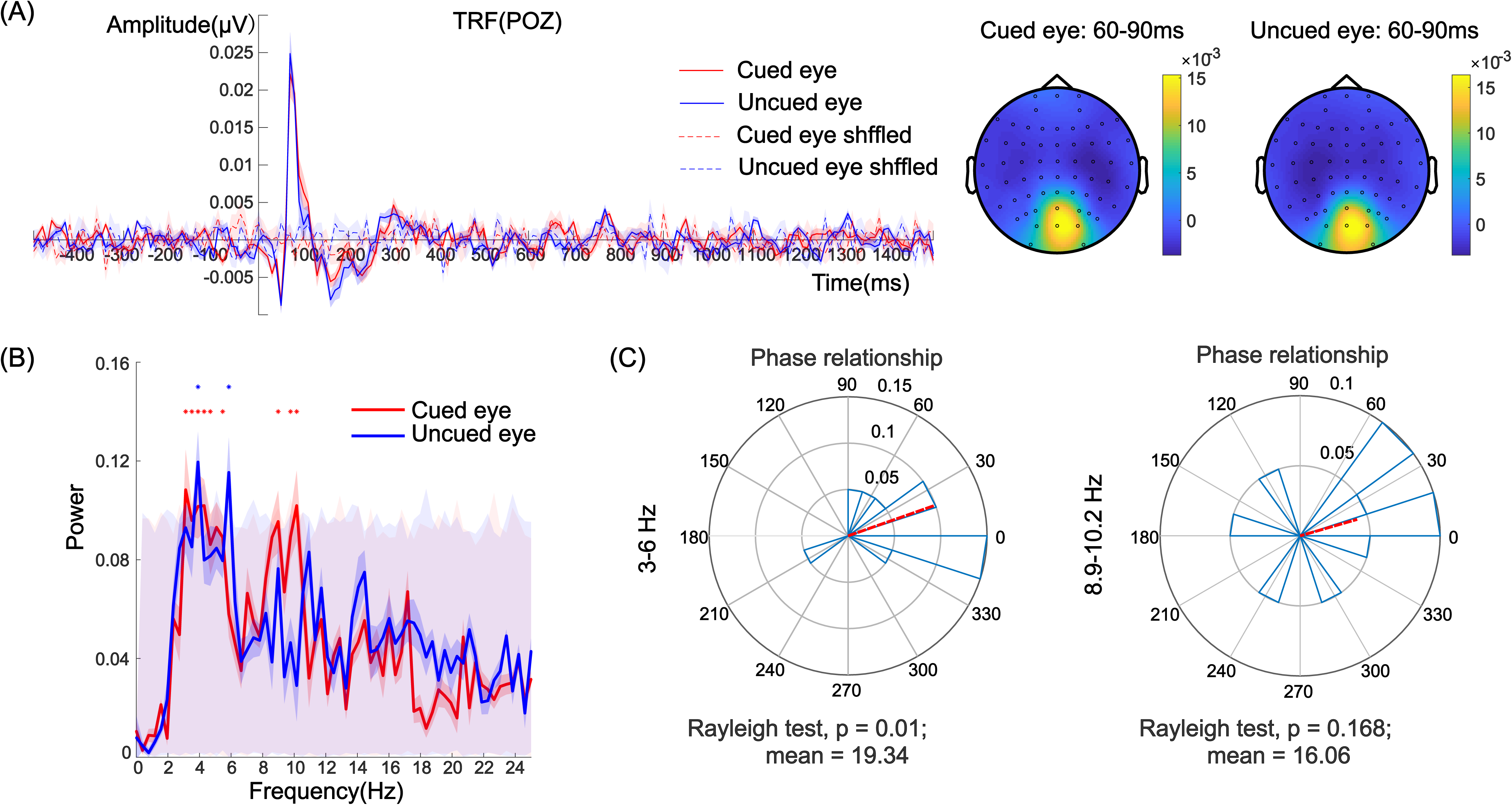
(A) TRF responses and topographic map of the two eyes at electrode POz. (B) Frequency map of the TRF waveform of the two eyes. The asterisk represents significant frequency points after the permutation test under each condition. The gray shaded area with red low contrast represents the frequency range of 1000 pseudo-TRFs. (C) Phase relationship with 3-6 Hz (theta band) and 8.9-10.2 Hz (alpha band) neural activity.

By combining the frequency and phase results of the TRFs at the Oz and POz electrodes, presenting cues in two eyes simultaneously can be shown to induce obvious attentional rhythms. Specifically, there was an in-phase mode between the two TRF responses, further supporting that the in-phase oscillation patterns observed in Experiment 1 indeed arise from the neural activity of binocular cells, which is consistent with our hypothesis. A one-way analysis of variance (ANOVA) was used to compare the phase differences between Experiment 1 and Experiment 2. We found that at electrode Oz, there were no significant phase differences in the band range of 2.7-6.3 Hz, *F*(1,18) = 0.47, *p* = 0.50, and the band range of 8.5-11 Hz, *F*(1.18) = 0.33, *p* = 0.58; in addition, at electrode POz, there were no significant phase differences in the band range of 2.7-6.3 Hz, *F*(1,18) = 0.10, *p* = 0.76, and in the band range of 8.5-11 Hz, *F*(1,18) = 1.31, *p* = 0.27. These findings further support the results and conclusions of Experiment 1.

## 4 General discussion

In this study, in combination with the use of temporal response tracking technology, we presented flashing discs in the central visual field of the left and right eyes to examine the neural activities induced by stimuli detected by the left and right eyes. First, we found that attentional rhythms were manifested by alpha and theta bands in the two experiments. Second, regardless of whether the cues were presented in one eye (Experiment 1) or both eyes (Experiment 2), the interaction pattern between the TRFs of each eye showed no significant differences and was always in phase, indicating that it was difficult for the cue stimulus to interfere with attentional rhythms in one eye alone. Thus, attention scarcely oscillated between the two eyes, implying that the neural sites that produce attentional rhythms are located in binocular cells.

Our results provide important neural evidence of attentional rhythms consistent with our previous behavioral oscillation research (Chen et al., 2018), further confirming the role of binocular cells in attentional rhythms. In a previous study, we used a high-resolution cue-target paradigm and found that regardless of whether the cue and the target were presented in the same eye or different eyes, the significant frequencies of behavioral oscillations were the same, proving that attentional oscillations originate from binocular cells (Chen et al., 2018). In behavioral studies, researchers always hope that the participants’ judgment criteria remain constant and that the participants maintain a stable state of attention in a high temporal resolution cue-target paradigm throughout the experiment. Only in this way can the researchers examine and analyze the periodic fluctuations in the participants’ accuracy data. Any changes in the participants’ attention state (such as sleepiness, distraction, fatigue, overexcitement, etc.) can undermine the measurement of behavioral oscillations and confound the results. However, the task of the high-resolution cue-target paradigm usually contains nearly 2000 trials and needs to be completed in several days or sessions (Chen, Tang, et al., 2017; Chen, Wang, et al., 2017; Fiebelkorn et al., 2013; Landau & Fries, 2012). It was difficult for the participants to maintain an ideal attention state throughout the experiment. More importantly, due to the limited trials of behavioral studies, researchers can only investigate low-frequency behavioral oscillations (VanRullen, 2016), which is another inherent difficulty in research investigating behavioral oscillations. Notably, compared to high-temporal behavioral methods, TRF is not only less susceptible to the influence of participants’ judgment criteria and attention state but also has two other advantages. First, TRF does not rely on participants’ behavioral responses, and the targets in the experiment were set to maintain participants’ attention focus on the flashing disc. Therefore, target detection accuracy is not used to analyze attentional rhythms and, thus, has no effect on the TRF responses. Second, instead of the 10 Hz frequency always used in SSVEP technology, the flicker frequency of the middle disc stimulation is 60 Hz in TRF technology, which is far beyond the critical frequency of human flash fusion (Wandell, 1995); thus, the flicker frequency and flicker patterns of the discs are subconscious information from the perspective of the participants, and it is difficult to blur attentional rhythms. More importantly, this study eliminated the possible interference (i.e., oscillations from switching between the visual fields) in Chen et al. (2018), resulting in a stricter investigation, and confirmed the role of monocular and binocular cells in attentional rhythms.

Furthermore, our results imply that the neural sites of attentional rhythms are located in the binocular cell visual pathway, which is consistent with the findings of previous studies. The phases of the theta band on occipital electrodes recorded with intracranial electrodes in epilepsy patients predict fluctuations in accuracy and reaction time (Helfrich et al., 2018). The theta band of neural oscillations in awake monkeys’ V4 cortex has also been shown to be the neural basis of 4 Hz behavioral oscillations (Kienitz et al., 2018). Relevantly, theta oscillations and theta band synchronization in the V1-V2 and V4-TEO regions are closely related to attentional rhythms (Spyropoulos, Bosman, & Fries, 2018). Cortical distance modulates attentional rhythms, and this modulation may be attributed to early cortical areas, such as V1, V2, and V3 (Chen et al., 2020). Using TRF, neural oscillations in the central posterior parietal lobe have been shown to be significantly related to behavioral oscillations (Jia et al., 2019). Notably, while monkeys performed an object-based attention task, Fiebelkorn et al. (2018) recorded their local field potentials and found that the phase of the theta band on the top frontal eye field (FEF) and lateral intraparietal area (LIP) area can predict the performance of behavioral oscillations (Fiebelkorn, Pinsk, & Kastner, 2018). Finally, importantly, in the above research, the occipital lobe, V4, posterior central parietal lobe, FEF, and LIP were found in the binocular cell visual pathway, which is consistent with the results of this study.

Compared with the large-scale interference in TMS studies, this study, which is based on the anatomical characteristics of the primary cortex, further found that attentional rhythms were produced in the binocular cell visual pathway instead of the monocular cell visual pathway.

## Acknowledgments

Thanks to Dr. Chencan Qian from the Institute of Biophysics, Chinese Academy of Sciences for the help in the production of spread spectrum stimulation. This work was supported by the National Natural Science Foundation of China (32100841, 32100842), MOE Project of Humanities and Social Sciences (20YJC190002), the Project of Social Science Foundation of Jiangsu Province (20YJC008), the Natural Science Foundation of Jiangsu Province (BK20190936), the Japan Society for the Promotion of Science KAKENHI (20K04381), the Project of Philosophy and Social Science Research in Colleges and Universities in Jiangsu Province (2019SJA1267).

## Notes

### Competing Interest Statement

The authors have declared no competing interest.

## References

Anstis, S., Verstraten, F. A., & Mather, G. (1998). The motion aftereffect. Trends in Cognitive Sciences, 2(3), 111–117.

Blakemore, C., & Campbell, F. W. (1969). On the existence of neurones in the human visual system selectively sensitive to the orientation and size of retinal images. The Journal of physiology, 203(1), 237–260.

Brainard, D. H. (1997). The Psychophysics Toolbox. Spatial Vision, 10(4), 433–436. doi:10.1163/156856897X00357

Busch, N. A., & VanRullen, R. (2010). Spontaneous EEG oscillations reveal periodic sampling of visual attention. Proceedings of the National Academy of Sciences of the United States of America, 107(37), 16048–16053. doi:10.1073/pnas.1004801107

Chen, A., Tang, X., Wang, A., & Zhang, M. (2017). Experimental paradigms for discrete attention in visual domain. Advances in Psychological Science, 25(6). doi:10.3724/sp.J.1042.2017.00923

Chen, A., Wang, A., Wang, T., Tang, X., & Zhang, M. (2017). Behavioral Oscillations in Visual Attention Modulated by Task Difficulty. Frontiers in Psychology, 8(1630). doi:10.3389/fpsyg.2017.01630

Chen, A., Wang, A., Wang, T., Tang, X., & Zhang, M. (2018). The primary visual cortex modulates attetion oscillation. Acta Psychologica Sinica, 50(2), 158–167.

Chen, A., Zu, G., Dong, B., & Zhang, M. (2020). Cortical Distance but Not Physical Distance Modulates Attentional Rhythms. Frontiers in Psychology, 11, 541085. doi:10.3389/fpsyg.2020.541085

Crosse, M. J., Di Liberto, G. M., Bednar, A., & Lalor, E. C. (2016). The multivariate temporal response function (mTRF) toolbox: A MATLAB toolbox for relating neural signals to continuous stimuli. Frontiers in Human Neuroscience, 10(604).

Dugué, L., Marque, P., & VanRullen, R. (2015). Theta oscillations modulate attentional search performance periodically. Journal of Cognitive Neuroscience, 27(5), 945–958. doi:10.1162/jocn_a_00755

Dugué, L., McLelland, D., Lajous, M., & VanRullen, R. (2015). Attention searches nonuniformly in space and in time. Proceedings of the National Academy of Sciences of the United States of America, 112(49), 15214–15219. doi:10.1073/pnas.1511331112

Dugué, L., Roberts, M., & Carrasco, M. (2016). Attention Reorients Periodically. Current Biology, 26(12), 1595–1601. doi:10.1016/j.cub.2016.04.046

Dugué, L., & Van Rullen, R. (2017). Transcranial magnetic stimulation reveals intrinsic perceptual and attentional rhythms. Frontiers in Neuroscience, 11(MAR). doi:10.3389/fnins.2017.00154

Dugué, L., Xue, A. M., & Carrasco, M. (2017). Distinct perceptual rhythms for feature and conjunction searches. Journal of Vision, 17(3). doi:10.1167/17.3.22

Fiebelkorn, I. C., & Kastner, S. (2019). A Rhythmic Theory of Attention. Trends Cogn Sci, 23(2), 87–101. doi:10.1016/j.tics.2018.11.009

Fiebelkorn, I. C., Pinsk, M. A., & Kastner, S. (2018). A Dynamic Interplay within the Frontoparietal Network Underlies Rhythmic Spatial Attention. Neuron, 99(4), 842–853 e848. doi:10.1016/j.neuron.2018.07.038

Fiebelkorn, I. C., Saalmann, Y. B., & Kastner, S. (2013). Rhythmic sampling within and between objects despite sustained attention at a cued location. Current Biology, 23(24), 2553–2558. doi:10.1016/j.cub.2013.10.063

Gilinsky, A. S., & Doherty, R. (1969). Interocular transfer of orientational effects. Science, 164(3878), 454–455.

Han, Q., & Luo, H. (2019). Visual crowding involves delayed frontoparietal response and enhanced top-down modulation. European Journal of Neuroscience, 50(6), 2931–2941. doi:10.1111/ejn.14401

Helfrich, R. F., Fiebelkorn, I. C., Szczepanski, S. M., Lin, J. J., Parvizi, J., Knight, R. T., & Kastner, S. (2018). Neural Mechanisms of Sustained Attention Are Rhythmic. Neuron, 99(4), 854–865.e855. doi:https://doi.org/10.1016/j.neuron.2018.07.032

Hembrook-Short, J. R., Mock, V. L., Martin Usrey, W., & Briggs, F. (2019). Attention enhances the efficacy of communication in V1 local circuits. Journal of Neuroscience, 39(6), 1066–1076. doi:10.1523/JNEUROSCI.2164-18.2018

Herbst, S. K., & Landau, A. N. (2016). Rhythms for cognition: the case of temporal processing. Current Opinion in Behavioral Sciences, 8, 85–93. doi:10.1016/j.cobeha.2016.01.014

Huang, Q., Jia, J., Han, Q., & Luo, H. (2018). Fast-backward replay of sequentially memorized items in humans. Elife, 7, e35164. doi:10.7554/eLife.35164.001

Hubel, D. H., & Wiesel, T. N. (1977). Ferrier lecture-Functional architecture of macaque monkey visual cortex. Proceedings of the Royal Society of London. Series B. Biological Sciences, 198(1130), 1–59.

Jensen, O., Bonnefond, M., & VanRullen, R. (2012). An oscillatory mechanism for prioritizing salient unattended stimuli. Trends in Cognitive Sciences, 16(4), 200–206. doi:10.1016/j.tics.2012.03.002

Jia, J., Fang, F., & Luo, H. (2019). Selective spatial attention involves two alpha-band components associated with distinct spatiotemporal and functional characteristics. NeuroImage, 199, 228–236. doi:https://doi.org/10.1016/j.neuroimage.2019.05.079

Jia, J., Liu, L., Fang, F., & Luo, H. (2017). Sequential sampling of visual objects during sustained attention. PLoS Biology, 15(6).

Kienitz, R., Schmiedt, J. T., Shapcott, K. A., Kouroupaki, K., Saunders, R. C., & Schmid, M. C. (2018). Theta Rhythmic Neuronal Activity and Reaction Times Arising from Cortical Receptive Field Interactions during Distributed Attention. Current Biology, 28(15), 2377–2387.e2375. doi:https://doi.org/10.1016/j.cub.2018.05.086

Klein, B. P., Fracasso, A., van Dijk, J. A., Paffen, C. L. E., te Pas, S. F., & Dumoulin, S. O. (2018). Cortical depth dependent population receptive field attraction by spatial attention in human V1. NeuroImage, 176, 301–312. doi:10.1016/j.neuroimage.2018.04.055

Lalor, E. C., Pearlmutter, B. A., Reilly, R. B., McDarby, G., & Foxe, J. J. (2006). The VESPA: A method for the rapid estimation of a visual evoked potential. NeuroImage, 32(4), 1549–1561. doi:10.1016/j.neuroimage.2006.05.054

Landau, A. N., & Fries, P. (2012). Attention samples stimuli rhythmically. Current Biology, 22(11), 1000–1004. doi:10.1016/j.cub.2012.03.054

Landau, A. N., Schreyer, H. M., Van Pelt, S., & Fries, P. (2015). Distributed Attention Is Implemented through Theta-Rhythmic Gamma Modulation. Current Biology, 25(17), 2332–2337. doi:10.1016/j.cub.2015.07.048

Livingstone, M. S., & Hubel, D. H. (1987). Psychophysical evidence for separate channels for the perception of form, color, movement, and depth. Journal of Neuroscience, 7(11), 3416–3468.

McCollough, C. (1965). Color adaptation of edge-detectors in the human visual system. Science, 149(3688), 1115–1116.

Mo, C., Lu, J., Wu, B., Jia, J., Luo, H., & Fang, F. (2019). Competing rhythmic neural representations of orientations during concurrent attention to multiple orientation features. Nature Communications, 10(1), 5264. doi:10.1038/s41467-019-13282-3

Motter, B. C. (1993). Focal attention produces spatially selective processing in visual cortical areas V1, V2, and V4 in the presence of competing stimuli. Journal of Neurophysiology, 70(3), 909–919. doi:10.1152/jn.1993.70.3.909

Paradiso, M. A., Shimojo, S., & Nakayama, K. (1989). Subjective contours, tilt aftereffects, and visual cortical organization. Vision Research, 29(9), 1205–1213.

Pelli, D. G. (1997). The VideoToolbox software for visual psychophysics: Transforming numbers into movies. Spatial Vision, 10(4), 437–442.

Re, D., Inbar, M., Richter, C. G., & Landau, A. N. (2019). Feature-Based Attention Samples Stimuli Rhythmically. Current Biology, 29(4), 693–699. doi:10.1016/j.cub.2019.01.010

Schoups, A. A., & Orban, G. A. (1996). Interocular transfer in perceptual learning of a pop-out discrimination task. Proceedings of the National Academy of Sciences, 93(14), 7358–7362.

Song, K., Meng, M., Chen, L., Zhou, K., & Luo, H. (2014). Behavioral Oscillations in Attention: Rhythmic α Pulses Mediated through θ Band. The Journal of Neuroscience, 34(14), 4837. doi:10.1523/JNEUROSCI.4856-13.2014

Spyropoulos, G., Bosman, C. A., & Fries, P. (2018). A theta rhythm in macaque visual cortex and its attentional modulation. Proceedings of the National Academy of Sciences of the United States of America, 115(24), E5614–E5623. doi:10.1073/pnas.1719433115

Su, Z., Wang, L., Kang, G., & Zhou, X. (2021). Reward makes the rhythmic sampling of spatial attention emerge earlier. Attention Perception Psychophysics, 83(4), 1522–1537. doi:10.3758/s13414-020-02226-5

Vanrullen, R. (2013). Visual attention: A rhythmic process? Current Biology, 23(24), R1110–R1112. doi:10.1016/j.cub.2013.11.006

VanRullen, R. (2016). Perceptual Cycles. Trends in Cognitive Sciences, 20(10), 723–735. doi:10.1016/j.tics.2016.07.006

VanRullen, R. (2018). Attention Cycles. Neuron, 99(4), 632–634. doi:10.1016/j.neuron.2018.08.006

Vanrullen, R., & MacDonald, J. S. P. (2012). Perceptual echoes at 10 Hz in the human brain. Current Biology, 22(11), 995–999. doi:10.1016/j.cub.2012.03.050

VanRullen, R., Reddy, L., & Koch, C. (2005). Attention-driven discrete sampling of motion perception. Proceedings of the National Academy of Sciences of the United States of America, 102(14), 5291–5296. doi:10.1073/pnas.0409172102

Wandell, B. A. (1995). Fundation of vision: Sunderland, Massachusetts, USA: Sinauer Associates.

Wassermann, E. M., Epstein, C. M., Ziemann, U., Walsh, V., Paus, T., & Lisanby, S. H. (2008). The Oxford Handbook of Transcranial Stimulation: New York, USA: Oxford University Press.

Zhang, H., Morrone, M. C., & Alais, D. (2019). Behavioural oscillations in visual orientation discrimination reveal distinct modulation rates for both sensitivity and response bias. Scientific Reports, 9(1), 1–11.

Zhaoping, L. (2008). Attention capture by eye of origin singletons even without awareness—A hallmark of a bottom-up saliency map in the primary visual cortex. Journal of Vision, 8(5), 1–1.

